# The S-palmitoylome and DHHC-PAT interactome of *Drosophila melanogaster* S2R+ cells indicate a high degree of conservation to mammalian palmitoylomes

**DOI:** 10.1101/2021.12.06.471463

**Authors:** Elena Porcellato, Juan Carlos Gonzalez, Constantin Ahlmann-Eltze, Mahmoud Ali Elsakka, Itamar Shapira, Jürgen Fritsch, Juan Antonio Navarro, Simon Anders, Robert B. Russell, Felix T. Wieland, Christoph Metzendorf

## Abstract

Protein S-palmitoylation, the addition of a long-chain fatty acid to target proteins, is among the most frequent reversible protein modification in Metazoa, affecting subcellular protein localization, trafficking and protein-protein interactions. S-palmitoylated proteins are abundant in the neuronal system and are associated with neuronal diseases and cancer. Despite the importance of this post-translational modification, it has not been thoroughly studied in the model organism *Drosophila melanogaster*.

Here we present the palmitoylome of Drosophila S2R+ cells, comprising 198 proteins, an estimated 3.5% of expressed genes in these cells. Comparison of orthologs between mammals and Drosophila suggests that S-palmitoylated proteins are more conserved between these distant phyla than non-S-palmitoylated proteins. To identify putative client proteins and interaction partners of the DHHC family of protein acyl-transferases (PATs) we established DHHC-BioID, a proximity biotinylation-based method. In S2R+ cells ectopic expression of the DHHC-PAT dHip14-BioID in combination with Snap24 or an interaction-deficient Snap24- mutant, as a negative control, resulted in biotinylation of Snap24 but not the Snap24-mutant. DHHC-BioID in S2R+ cells using 10 different DHHC-PATs as bait identified 520 putative DHHC-PAT interaction partners of which 48 were S-palmitoylated and are therefore putative DHHC-PAT client proteins. Comparison of putative client protein/DHHC-PAT combinations indicates that CG8314, CG5196, CG5880 and Patsas have a preference for transmembrane proteins, while S-palmitoylated proteins with the Hip14-interaction motif are most enriched by DHHC-BioID variants of approximated and dHip14. Finally, we show that BioID is active in larval and adult Drosophila and that dHip14-BioID rescues dHip14 mutant flies, indicating that DHHC-BioID is non-toxic.

In summary we provide the first systematic analysis of a Drosophila palmitoylome. We show that DHHC-BioID is sensitive and specific enough to identify DHHC-PAT client proteins and provide DHHC-PAT assignment for ca. 25% of the S2R+ cell palmitoylome, providing a valuable resource. In addition, we establish DHHC-BioID as a useful concept for the identification of tissue-specific DHHC-PAT interactomes in Drosophila.

## 1. Introduction

Protein S-acylation, the attachment of a fatty acid to a cysteine residue of a protein via a thioester bond, is one of the most abundant reversible post-translational modifications in Metazoa [1, 2]. Because the fatty acid palmitate is most frequently involved [3–5], protein S-acylation is often referred to as S-palmitoylation (used here) or just palmitoylation.

Though the functional effects of S-palmitoylation on target proteins are often specific, some common functions of S-palmitoylation have been identified. For example, S-palmitoylation of soluble proteins results either in the reversible regulation of their subcellular localization and trafficking or their tight membrane association (reviewed in [6]). In the case of transmembrane proteins S-palmitoylation can affect trafficking, localization to specific membrane domains, complex formation with other proteins, interplay with other post-translational modifications and initiation of conformational changes ([7] and reviewed in [8]). Thus, the effects of S-palmitoylation on target proteins are very diverse; the biological functions of target proteins include iron uptake (transferrin receptor) [9], ER quality control/ER Ca^2+^ homeostasis (calnexin) [10] and integrin signaling [11]. Several S-palmitoylated proteins are associated with diseases such as cancer and neurological diseases [1], suggesting that a better understanding of protein S-palmitoylation and its effects on target protein function is of interest to biomedical research.

Most protein S-palmitoylation reactions are catalyzed by the DHHC protein S-acyltransferase family (DHHC-PATs) with the reverse process (de-acylation) being performed by thioesterases. DHHC-PAT enzymes are characterized by four to six transmembrane domains and the name-giving DHHC (Asp-His-His-Cys) motif in an approximately 50-residue-long cysteine-rich domain (CRD) [12]. The *Saccharomyces cerevisiae* genome codes for seven DHHC-PATs, 15 are found in *Caenorhabditis elegans*, 24 in most mammals and 22 in *Drosophila melanogaster* [12–14]. De-acylation is carried out by four thioesterases in mammals (palmitoyl-protein thioesterase 1 and 2 (PPT1 and PPT2, respectively) and acyl-protein thioesterase 1 and 2 (APT1 and APT2) and by three thioesterases in Drosophila (Ppt1, Ppt2 and Apt1 [14]). While PPT1 and PPT2 act in lysosomes to remove acyl chains from client proteins [15, 16], APT1 and APT2 are responsible for the dynamic S-palmitoylation/de-palmitoylation cycle at the cytoplasmic membrane [17]. In both, mammals and insects, the different thioesterases have substrate specificity and cannot substitute each other [14,18,19].

Although mammals and insects have approximately the same number of DHHC-PATs, there are several DHHC-PATs lacking clear orthology between the two clades [14]. Nine Drosophila DHHC-PAT genes are exclusively expressed in the adult testis, and five of these have a genomic arrangement indicating recent duplication events [14]. Seven of the other 13 DHHC-PATs are most abundantly expressed in nervous tissue [14]. Hence, S-palmitoylation seems to play important roles in neurons and the male reproductive organ of Drosophila. In mammals, the highest fraction of S-palmitoylated proteins is found in neuronal tissues [1]. Neither DHHC-PATs expressed specifically in the testis, nor a testis palmitoylome has been reported in mammals. It is unclear whether this is due to a lack of research or an indication that S-palmitoylation does not play an important role in the mammalian testis.

Despite the many known S-palmitoylated proteins in mammals (>5000 proteins [1, 2]), not much is known in invertebrate model organisms [20]. In Drosophila, protein S-palmitoylation has mostly been studied in the context of specific proteins. For example, it was reported that the expression of the Drosophila ortholog of the DHHC-PAT Huntingtin-interacting protein 14 (dHip14) is required in the nervous system for survival to adulthood and that it interacts with SNAP25 and cysteine string protein (CSP) [21, 22]. dHip14 function is also required for S-palmitoylation and secretion of the short gastrulation (Sog) protein [23]. The DHHC-PAT approximated (app), which is most similar to human DHHC-PATs DHHC9/DHHC14/DHHC18, was shown to have DHHC-PAT-dependent and independent functions. app binds to and localizes Dachs to the apical junctional region of imaginal discs and palmitoylates the large protocadherin Fat, resulting in repression of Fat function and promoting tissue growth [24, 25]. Finally, RNAi-mediated knockdown of CG5196, an ER-localized DHHC-PAT similar to human DHHC6, results in more (and smaller) units of Golgi-ER exit sites as well as an increased cell size phenotype [26]. A more comprehensive catalogue of S-palmitoylated proteins in Drosophila and its systematic analysis is so far lacking, but would provide a valuable resource for the community. Furthermore, tools to identify client proteins of DHHC-PATs on the proteomic scale and in specific tissues are yet to be developed.

To this end, we herein provide a detailed study of the palmitoylome of S2R+ cells and use a novel, BioID-based approach to identify potential DHHC-PAT interaction partners and putative client proteins of DHHC-PATs (proteins that are palmitoylated by a given DHHC-PAT).

## 2. Methods

### 2.1 Molecular cloning

pENTR-DHHC plasmids were kindly provided by C. Korey [14]. The following plasmids were obtained from AddGene (AddGene plasmid IDs in parentheses): pcDNA-mcs-BioID2-HA (#74224), pcDNA-Myc-BioID2-mcs (#74223), pcDNA3_1_MCS-BirA(R118G)-HA (#36047), pcDNA3_1_Myc-BirA(R118G)-MCS (#35700), pGWvectorBirA(R118G)-HA (#53581) and pMT-puro (#17923). pUASTattB (GeneBank accession numer EF362409.1) [27] was obtained from Johannes Bischof, FlyORF, Zürich, Switzerland. pMT-GAL4 was obtained from the Sinning lab (BZH, Heidelberg University, Germany). Restriction endonucleases were obtained from New England BioLabs, Phusion polymerase for PCR and T4 ligase for DNA-ligation were from Thermo Scientific. Unless stated otherwise, CaCl_2_ competent DH5alpha were used for transformation. Primers were obtained from Biomers.net, Germany, and all clones were validated by Sanger sequencing (GATC Biotech, Germany).

pUASTattB-mycBioID and pUASTattB-mycBioID2 were generated by amplifying NotI-mycBioID-XhoI and NotI-mycBioID2-XhoI from pcDNA3_1_Myc-BirA(R118G)-MCS and pcDNA-Myc-BioID2-mcs, respectively. pUASTattB and PCR-products were NotI/XhoI digested and products ligated accordingly.

pMT-GW-BioID-HA-puro was generated by digesting pGWvectorBirA(R118G)-HA with StuI and PmeI to liberate the Gateway Cassette (Invitrogen) together with the BioID-HA fragment. pMT-puro was linearized by digestion with EcoRV and ligated with the pGWvecorBirA(R118G)-HA digestion products. Ligation products were transformed into ccdB-survival bacteria (Invitrogen). Clones harboring pMT-GW-BioID-HA-puro were identified by differential antibiotic selection and validated by sequencing.

pMT-DHHC-BioID-HA vectors were generated through LR-clonase-mediated recombination (Gateway® Technology, Invitrogen) of pENTR-DHHC plasmids [14] with pMT-GW-BioID-HA-puro as destination vector according to the manufacturer’s protocol.

pUASTattB-dHip14-BioID-HA was generated by Enhanced Gibson Assembly (New England BioLabs) of a dHip14 5′UTR PCR product, a dHip14-BioID-HA PCR product, a dHip14 3′UTR PCR product and linearized pUASTattB. The 5′UTR and the 3′UTR of *dHip14* were PCR amplified from genomic DNA and dHip14-BioID-HA was PCR amplified from pMT-dHip14-BioID-HA-puro using primers specified in Supplementary Table S1.

pMT-FLAG-dSnap24, pMT-FLAG-Snap25-PA and pMT-FLAG-dCSP were generated by cloning the respective coding regions from cDNA (total RNA from flies or S2R+ cells reverse transcribed with RevertAid First Strand cDNA Synthesis Kit (Thermo-Fisher) according to the manufacturer’s protocol), using Xho-FLAG-flanked forward primers and NotI-flanked reverse primers, respectively (Supplementary Table S1), and then ligated in XhoI/NotI-digested pMT-puro.

### 2.2 Mutagenesis of dSnap24 and dSnap25

Site-directed mutagenesis was performed using pMT-FLAG-SNAP24 (SNAP24 P120A) and pMT-FLAG-SNAP25 (SNAP25 P125A) as templates for amplification with PfuPlus DNA polymerase (Roboklon) in combination with the respective primers (Supplementary Table S1). Primers for mutagenesis were generated with the help of the Agilent Quickchange design tool. Parental DNA was digested via incubation with DpnI (New England BioLabs) before transforming the amplification products into CaCl_2_ competent DH5alpha. Mutant plasmids were validated by Sanger sequencing.

### 2.3 Cell culture

The Drosophila S2R+ cell line [28] was obtained from the Teleman lab (DKFZ, Heidelberg, Germany) and was cultured at 25 °C in Drosophila Schneider’s medium (Invitrogen) supplemented with 10% heat-inactivated fetal bovine serum (Gibco) and 1X (100 U/mL) penicillin/streptomycin (Gibco), unless indicated otherwise.

For transient transfection cells were seeded and after 18 h cells were transfected with plasmid DNA using X-tremeGENE HP DNA transfection reagent (Roche), according to the manufacturer’s protocol (for 12-well plate format: 800 μL of 500 000 cells/mL; 500 ng plasmid, ratio of plasmid to reagent 1:8).

To select for stable transfected cells, cells were cultured in selective medium containing 1 μg/μL Puromycin (Invitrogen) from approximately 5 days after transfection, depending on cell density.

Protein expression via the metallothionine promoter was induced by supplementing the growth medium with 0.25 mM CuSO_4_ (Sigma). For BioID experiments, medium was further supplemented with 50 μM biotin (Sigma), unless indicated otherwise.

Transfected cells were harvested as follows, unless specified otherwise. 24–72 h after induction of protein expression cells were washed with phosphate buffered saline (PBS), scraped in a small volume of PBS, pelleted (6,000 x *g*, 5 min, 4 °C), resuspended in ice-cold RIPA buffer (50 mM Tris-HCl, pH 7.4, 1% NP-40, 0.25% Na-deoxycholate, 150 mM NaCl; 1 mM EDTA with protease inhibitor cocktail) and sonicated for 5 min in an ice-cold sonication bath and centrifuged (10 000 x *g*, 5 min, 4 °C) to remove insoluble debris. The supernatant was used immediately or stored at -80 °C.

### 2.4 Fly maintenance and generation of transgenic flies

Flies were maintained on sucrose instant mashed potato food (13 g/L agar, 49 g/L sucrose, 36 g/L instant mashed potato powder (demeter bio quality), 17 g/L dry yeast, 7.95 mL/L 10% Nipagen and 2.16 mL/L propionic acid) under a 12 h / 12 h light/dark cycle at 25 °C unless stated otherwise. CO_2_ delivered through a CO_2_-pad stage was used for anesthesia for fly sorting.

Biotin supplementation of fly food was achieved by adding the appropriate amount of 164 mM biotin in DMSO to fly food cooled to approximately 45 °C, followed by thorough mixing. *w^1118^* and *w^-^; elavGAL4/TM6B, Tb* were from Juan Navarro (Regensburg University, Germany), *w^-^; elavGAL4 / CyO* (Bloomingon #8765) was from Bloomington Stock center, *w^-^; +; daGAL4* was from Maria Lind Karlberg (Uppsala University, Sweden), *w^-^; If/CyO; neoFRT80B, Hip14^2^/TM6B, Tb, Sb* was derived from Bloomington #39734 [22], *w^-^; If/CyO; MKRS, Sb/TM6B, Tb, Hu* was from Norman Zielke (ZMBH, Heidelberg University, Germany) and flies with the *dHip14* alleles *FRT79, dHip14^ex11^* and *dHip14^ex12^* were from Steven Stowers (Montana State University, USA). *UAS-Dnz1-3xHA* flies were obtained from FlyORF [29]. Fly lines *w^-^; Hip14^ex11^ / TM6B, Tb, Hu* and *w^-^; elavGAL4 / CyO; Hip14^ex12^ / TM6B, Tb, Hu* and *w^-^; elavGAL4 / CyO; Hip14^2^ / TM6B, Tb, Hu* and *w^-^; UAS-Hip14-BioID / CyO; Hip14^2^ / TM6B, Tb, Hu* and *w^-^; If / CyO; UAS-Dnz1-3xHA, Hip14^ex12^ / TM6B, Tb, Hu* and *w^-^; If / CyO; UAS-Dnz1-3xHA, Hip14^ex12^ / TM6B, Tb, Hu* were generated by standard fly genetics through crossing and recombination.

Lines *w^1118^;+;UAST-mycBioID* and *w^1118^;+;UAST-mycBioID2* and *w^1118^; UAST-dHip14-BioID-HA* were generated by phi31-mediated site-directed integration [27, 30] by injection of plasmids pUASTattB-mycBioID and pUASTattb-mycBioID2 into fly line R8622 (Rainbow Transgenic Flies, Inc, Camarillo, California, USA) carrying the *phi31* gene on the X chromosome and *attP* on the third chromosome (3L68A4). pUASTattB-dHip14-BioID-HA was injected and integrated into fly line RB25709 (Rainbow Transgenic Flies, Inc.) with *phi31* on the X chromosome and *attP* on the second chromosome (25C6). Injections were done by Rainbow Transgenic Flies, inc, Camarillo, Californa, USA. phi31 integrase was removed by crossing male recombinant flies with female *w^1118^* for two generations followed by six generations of outcrossing female flies with *w^1118^* flies.

### 2.5 BioID in flies

For BioID experiments the respective parents were allowed to lay eggs on the indicated food source for 24-48 h at 25 °C. After further 48 h vials were transferred to 30 °C. To prepare samples for WB, larvae or adult animals, heads, thoraces or abdomen were homogenized in cold RIPA buffer (50-100 μL per L3 larvae or adult) using a glass homogenizer or for small sample volumes a plastic micro pestle.

### 2.6 Isolation of Biotinylated proteins

Biotinylated proteins from BioID experiments were purified for proteomic analysis with Dynabeads MyOne Streptavidin C1 (Invitrogen) as previously described [31]. Briefly, cell lysates in RIPA buffer (500 μg) were incubated overnight at 4 °C with 50 μL slurry of Dynabeads MyOne Streptavidin C1 (Invitrogen) and then washed for 10 min on a rotator at room temperature as follows: twice with washing buffer 1 (2% SDS in ddH_2_O), twice with washing buffer 2 (0.1% deoxycholate, 1% Triton X-100, 500 mM NaCl, 1 mM EDTA, and 50 mM HEPES, pH 7.5), twice with washing buffer 3 (250 mM LiCl, 0.5% NP-40, 0.5% deoxycholate, 1 mM EDTA, and 10 mM Tris, pH 8.1) and twice with washing buffer 4 (50 mM Tris, pH 7.4, and 50 mM NaCl). Biotinylated proteins were dissociated from beads by incubation with 25 μL elution buffer (10 mM Tris, pH 7.4, 2% SDS, 5% (v/v) β-mercaptoethanol and 2 mM biotin) for 15 min at 98 °C.

### 2.7 Acyl-RAC assay

Isolation of S-palmitoylated proteins was performed as previously described [32]. Briefly, S2R+ cells were washed in PBS, scraped, pelleted (2000 x *g*, 4 °C, 5 min) and resuspended in Buffer A (25 mM HEPES, 25 mM NaCl, 1 mM EDTA, protease inhibitor cocktail, pH 7.5). After sonication, cell debris was removed by centrifugation (800 x *g*, 4 °C, 10 min), membranes were enriched by centrifugation of the resulting supernatant at 16,200 x *g* for 50 min and 4°C. The pelleted membranes were lysed in Buffer A containing 0.5% Triton X-100.

Equal amounts of protein (1 mg for mass spectrometry) were diluted 1:2 with blocking buffer (100 mM HEPES, pH 7.5, 1 mM EDTA, 2.5% SDS, 2.5% MMTS) at 40 °C for 2 h. Proteins were precipitated at -20 °C for 20 min after addition of three volumes of ice cold acetone. The pellet was washed with 70% acetone, air dried and resuspended in 300 μL binding buffer (100 mM HEPES, 1.0 mM EDTA, 1% SDS, pH 7.5) followed by the addition of 0.03 g Thiopropyl Sepharose 6B beads (GE Healthcare) (washed with binding buffer). Either 2 M NH_2_OH, pH 7.5 (+HA samples) or 2 M Tris-HCl, pH 7.5 (-HA samples) was added to a final concentration of 0.5 M and proteins were allowed to bind at room temperature overnight on a rotator. Beads were washed 5 times with binding buffer before 20 μL 4x reducing Laemmli buffer were added. Bound proteins were recovered by incubation at 95 °C for 10 min followed by SDS-PAGE.

### 2.8 Protein identification by mass spectrometry

Eluted material from the beads was separated by 1D SDS-PAGE (NuPage 125 gels, Invitrogen). Separated proteins were visualized by colloidal Coomassie blue staining. The proteins were reduced with 60 µL 40 mM dithiothreitol (DTT; Sigma-Aldrich, Taufkirchen, Germany) in 50 mM TEAB, pH 8.5 at 57 °C for 30 min and alkylated with 60 µL 50 mM iodoacetamide (IAA; Sigma-Aldrich, Taufkirchen, Germany) in 50 mM TEAB, pH 8.5 at 25 °C for 20 min in the dark. Gel pieces were dehydrated with 60 µL 100% ACN and washed with 60 µL 50 mM TEAB, pH 8.5. A total of 30 µL 8 ng/µL in 50 mM TEAB trypsin solution (sequencing grade, Thermo-Fisher, Rockford, USA) was added to the dry gel pieces and incubated for 6 h at 37 °C. The reaction was quenched by the addition of 20 µL 0.1% trifluoroacetic acid (TFA; Biosolve, Valkenswaard, Netherlands). The resulting peptides were extracted by two dehydration steps in 20 µL ACN for 20 min each followed by washing in 30 µL 50 mM TEAB, pH 8.5. The supernatant from each extraction step was collected and dried in a vacuum concentrator before LC-MS analysis. Samples were diluted in 15 μL 0.1% TFA, 99.9% water. Nanoflow LC-MS^2^ analysis was performed with an Ultimate 3000 liquid chromatography system directly coupled to an Orbitrap Elite mass spectrometer (both Thermo Fisher, Bremen, Germany). Samples were delivered to an in-house packed analytical column (inner diameter 75 µm x 20 cm; CS – Chromatographie Service GmbH, Langerwehe, Germany) filled with 1.9 µm ReprosilPur-AQ 120 C18 material (Dr. Maisch, Ammerbuch-Entringen, Germany). Solvent A was 0.1% formic acid (FA; ProteoChem, Denver, CO, USA) and 1% ACN (Biosolve) in H_2_O (Bisolve). Solvent B was composed of 0.1% FA (ProteoChem), 10% H_2_O (Biosolve) and 89.9% ACN (Biosolve). Samples were loaded onto the analytical column for 20 min with 3% solvent B at a flow rate of 550 nL/min Peptide separation was carried out with a 100 min linear gradient (3-23% solvent B) and a 20 min linear gradient (23-38% solvent B) with a reduced flow rate of 300 nL/min. The mass spectrometer was operated in data-dependent acquisition mode, automatically switching between MS and MS^2^. MS spectra (m/z 400–1600) were acquired in the Orbitrap at 60 000 (m/z 400) resolution. Fragmentation in the HCD cell was performed for up to 15 precursors, and the MS^2^ spectra were acquired.

The raw mass spectrometry data were processed with MaxQuant [33]. The resulting protein intensities were normalized by the median difference between each sample and the negative controls. The log fold-changes and the respective significance of the change were calculated using proDA version 0.1.0 [34]. Unlike alternative methods that rely on imputation to handle missing values, proDA provides a principled statistical test and is better able to control the false discovery rate. proDA calculates a dropout curve for each protein intensity and combines it with empirical Bayesian priors to determine if the observed pattern of observed and missing values and the corresponding mean difference could be simply due to chance.

### 2.9 Bioinformatic data analysis

Protein identifiers, sequences, descriptions, and gene names were obtained from UniProt [35]. The DRSC Integrative Ortholog Prediction Tool (DIOPT) was used to determine mammalian orthologs for the Drosophila proteins by searching against human, mouse and rat proteomes and selecting only those orthologs with the highest confidence [36]. The SwissPalm database was used to determine the S-palmitoylation state of mammalian proteins [2, 37]. Mammalian proteins reported as S-palmitoylated by at least one targeted study, or by two different experimental techniques were additionally classified as high confidence. S-palmitoylation was predicted using the program CSS-Palm 4.0 with high threshold [38]. Functional enrichment analyses of Gene Ontology (GO) terms were performed using Fisher’s Exact test and False Discovery Rate (FDR) for multiple testing correction, through the PANTHER overrepresentation tool [39, 40].

### 2.10 Western blotting

Protein content in cell culture lysates and eluates from biotin/streptavidin or acyl-RAC mediated protein purification was determined by the BCA assay (Pierce, Thermofisher Scientific) and mixed with 4x reducing Laemmli buffer to yield the indicated protein amounts in 1x Laemmli buffer. Unless stated otherwise, samples were boiled for 5 min. Proteins were then separated by SDS-PAGE gel electrophoresis (BioRad) and blotted onto PVDF membranes. Antibodies were preparedin blocking buffer (PBS-T in 5% BSA) as follows: mouse polyclonal anti-myc (serum from the lab); mouse monoclonal anti-HA (1:10 000; Thermo-Fisher #26183); mouse or rabbit monoclonal anti-FLAG (1:10 000; Thermo-Fisher #14-6681-82 and #701629). Quantitative immunodetection of tagged proteins and biotinylated proteins was carried out using a LI-COR infrared imager after incubation with the following antibodies/reagents: Goat anti-Mouse IgG (1:10 000; LI-COR #925-68020 IRDye 680LT Goat anti-Mouse IgG); Goat anti-Rabbit IgG (1:10 000; LI-COR # 926-68021 IRDye 680LT Goat anti-Mouse IgG), fluorescent Streptavidin 1:10 000; LI-COR IRDye® 800CW Streptavidin.

### 2.11 Data Quantification and Statistics

Relative promiscuous biotinylation in myc-BioID experiments was estimated by integrating streptavidin signals in each lane normalized to the expression of BioID or BioID2. In co-overexpression experiments, relative biotinylation signals of FLAG-dSNAP24/25/25* and CSP detected via Western blot were quantified as follows: biotinylation raw signals were normalized to the relative expression of the correspondent DHHC-PAT-BioID and expression of the FLAG-tagged protein. To calculate fold-changes of biotinylation the normalized biotinylation signal of the sample was divided by the normalized biotinylation of the negative control sample (only DHHC-BioID expression but no FLAG-tagged protein expression). Ratios were then normalized by the values calculated in S2R+ cells not expressing any of the DHHC-PAT-BioID. One-way ANOVA (GraphPad Prism Version 5.0 for Windows/Mac OS, GraphPad Software, San Diego, California, USA) was used for statistical analyses.

## 3. Results and discussion

### Drosophila S2R+ cells express at least 198 S-palmitoylated proteins

Extensive proteomic data exist on the S-palmitoylation status of mammalian cell lines and tissues [1, 2], however, very little is known about the degree of protein S-palmitoylation in Drosophila. Here, we used acyl-resin assisted capture (acyl-RAC) [32] and LC-MS/MS to purify and identify potentially S-palmitoylated proteins from the membrane fraction of S2R+ cells (Supplementary Data File SDF1). To deal with missing data in negative controls, a result of affinity purification, we applied a novel method that uses the overall dropout probability for each intensity and empirical Bayesian priors to calculate a principled statistical test [34].

We initially identified 1188 different proteins by mass spectrometry (Supplementary Data File SDF1), then filtered for those significantly enriched by acyl-RAC (proteins with a fold change (FC) equal to or above 2 and a false discover rate (FDR) below 0.1), removed proteins lacking cysteine residues, and excluded six proteins considered to be false positives, e.g. enzymes with thioester bonds known to not play a role in S-palmitoylation in the sense of the here investigated reversible protein lipidation. This resulted in 198 proteins that we considered S-palmitoylated, of which 51 were highly enriched by acyl-RAC (FC >= 20) and thus additionally labelled as high confidence (HC) S-palmitoylated proteins (Figure 1A; Supplementary Table S2).

**Figure 1:**
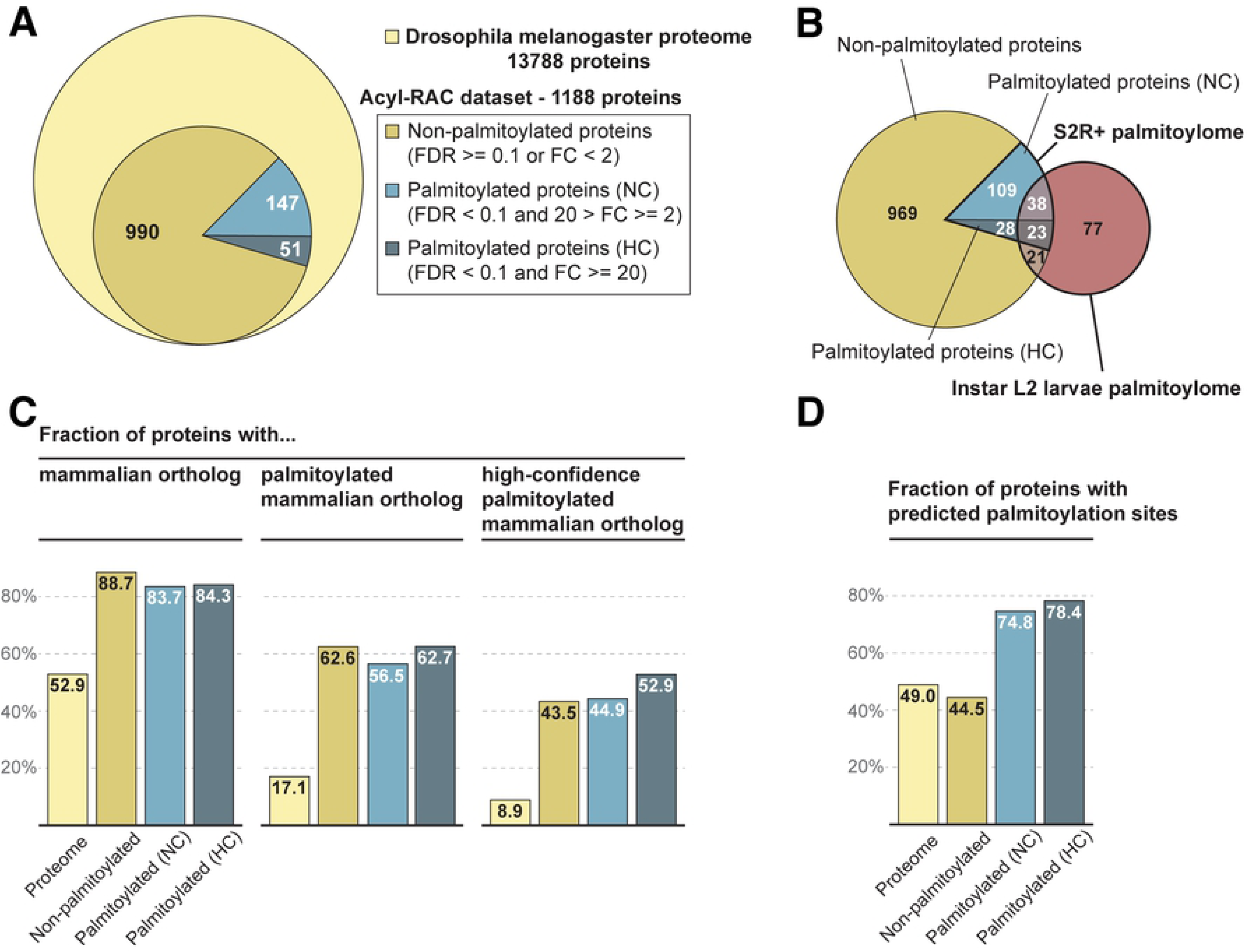
Drosophila S2R+ cell palmitoylome evaluation. (A) Acyl-RAC dataset subjected to different filtering to identify proteins that are unlikely to be palmitoylated (“non-palmitoylated”, FDR >= 0.1 or FC < 2), palmitoylated proteins with “normal” confidence (“NC”, FDR < 0.1 and FC < 20 and FC >= 2), palmitoylated proteins with “high” confidence (“HC”, FDR <0.1 and FC >= 20). (B) Overlap of our acyl-RAC dataset, consisting of the S2R+ palmitoylome proteins (normal and high confidence) and proteins not considered to be palmitoylated, with the larval palmitoylome from Strassburger et al 2019 (Strassburger et al. 2019). (C) Barplots indicate the fraction (%) of proteins that have a mammalian ortholog (left), that have a mammalian ortholog that is palmitoylated (center) and that have a mammalian ortholog that is palmitoylated with “high” confidence (right) in each of the protein sets defined above. The palmitoylation status of the mammalian orthologs was obtained from the SwissPalm database. (D) Barplot that shows the fraction (%) of proteins predicted to be palmitoylated by CSS-Palm in each of the protein sets.

Next, we compared our S2R+ cell palmitoylome to that of Drosophila instar L2 larvae [20]. Of the 159 S-palmitoylated proteins in larvae, 61 overlapped with 30% of the 198 S-palmitoylated proteins in S2R+ cells. 21 larval S-palmitoylated proteins overlapped with 2% of the 990 proteins that were not significantly S-palmitoylated in S2R+ cells (Figure 1B). This comparison may indicate that only a very small fraction of proteins that are not significantly S-palmitoylated in S2R+ cells are likely to be S-palmitoylated in another tissue. However, the identification of more S-palmitoylated proteins in S2R+ cells (198 proteins) than in larvae (159 proteins) was unexpected. Whole larvae are more complex than S2R+ cells and would be expected to contain a more diverse palmitoylome than the cell line. The difference is most likely due to technical reasons; we used acyl-RAC, whereas Strassburger *et al.* used acyl/biotin exchange (ABE) [41] for enrichment and purification of S-palmitoylated proteins. As ABE requires many precipitation and resolubilization steps it is more prone to loss of proteins than acyl-RAC. Also the data analysis strategies were different; we used MaxQuant for protein identification and label-free quantification while MuDPIT [41] was used by [20], which may further contribute to differences in peptide detection, protein identification and protein quantification. Finally, a mix of different tissues, as in whole larval homogenates, may result in the dilution of tissue-specific S-palmitoylated proteins below the limit of detection. Therefore, tissue and cell-type-specific palmitoylomes may favor the recovery of more complete palmitoylomes compared to whole-organism approaches.

In conclusion, we identified 198 S-palmitoylated proteins, of which we classified 51 as high confidence S-palmitoylated proteins. Furthermore, 61 S-palmitoylated proteins identified by both studies (larvae and S2R+ cells) represent the currently best-supported set of S-palmitoylated proteins in Drosophila (Supplementary Table S3).

### Palmitoylated proteins are overrepresented in orthologs of Drosophila and mammals

We used multiple approaches to assess the degree of conservation in protein S-palmitoylation between Drosophila and mammals. While the fraction of genes encoding for S-palmitoylated proteins in mammals is approximately 10-14% [1] our initial estimation in Drosophila is clearly lower: if we consider that S2R+ cells express 5885 genes on average as reported previously [42], the 198 proteins we determined as being S-palmitoylated represent only 3.4%. However, because technical limitations in proteomics are greater than for transcriptomics, this is likely an underestimate. Also, S2R+ cells may not be the cell type with the most diverse palmitoylome; in Drosophila, neuronal tissue and testes may yield higher numbers of S-palmitoylated proteins as the expression of several DHHC-PATs is most abundant in these tissues [14]. To estimate the percentage of palmitoylated proteins on the whole-body level in Drosophila, we determined the fraction of Drosophila proteins with S-palmitoylated mammalian orthologs and all Drosophila orthologs of mammalian proteins (Figure 1C; Supplementary Table S2). This number, 17.1% is clearly higher than 10-14% palmitoylated proteins in the full mammalian proteome. That S-palmitoylated proteins are enriched within the group of Drosophila/mammalian orthologs indicates a higher degree of evolutionary conservation of S-palmitoylated proteins. This is expected, as S-palmitoylated proteins perform many essential cellular functions [43, 44]. However, 17.1% could also be an overestimation of the number of palmitoylated proteins in Drosophila, as not each palmitoylated protein in mammals need to be palmitoylated in Drosophila. However, to determine the overlap of palmitoylated proteins between mammalian and Drosophila will require a much more detailed Drosophila palmitoylome that covers similar tissues as covered by the mammalian palmitoylomes.

We did similar ortholog searches for the proteins in the three subsets from our acyl-RAC experiment, as defined above (palmitoylated, palmitoylated with high confidence, and non-palmitoylated). As expected, the percentage of proteins with S-palmitoylated mammalian orthologs is much higher within the S2R+ palmitoylome (56.5% and 62.7%, in the palmitoylome sets with normal and high confidence, respectively) than in the complete proteome, thus supporting the validity of our experiment to recover real palmitoylated proteins (Figure 1C). Interestingly, 62.6% of non-palmitoylated proteins from the acyl-RAC dataset were also found to have mammalian S-palmitoylated orthologs; this could be an indication that there are likely proteins that are less frequently palmitoylated in S2R+ cells than in other cell types and tissues.

Overall, we suggest that the proportion of S-palmitoylated proteins in Drosophila is similar to that in mammals (10-14%) as the current proportion estimate ranges from 3.5% between 17.1%.

### Prediction of protein S-palmitoylation in Drosophila using CSS-Palm may yield a high false-positive rate

As S-palmitoylation prediction programs are trained on available data that are biased towards mammals and currently lack sites from insects, it is not clear how faithful they will be in Drosophila. However, the high apparent conservation between Drosophila and mammalian S-palmitoylated proteins suggests that they might nevertheless be applicable to insects. We thus applied CSS-Palm [38] to three groups within the acyl-RAC dataset: non-palmitoylated, S-palmitoylated (NC) and S-palmitoylated (HC), and in the Drosophila reference proteome for comparison. Surprisingly, 49% of proteins in the Drosophila reference proteome have at least one predicted site (Figure 1D), which is more than 10 times higher than the percentage we extrapolated from our palmitoylome dataset (3.5%), and 3 times higher than when using mammalian orthologs as a reference (17.1%). Within the acyl-RAC dataset, 44.5% of the 990 non-palmitoylated proteins and about 75% (74.8% and 78.4% of the normal confidence and high confidence S-palmitoylated proteins, respectively) are predicted to be S-palmitoylated. Hence, CSS-Palm seems to be able to predict these proteins in Drosophila. However, because a very high number of proteins predicted have very low experimental support (low FC in acyl-RAC) compared to a low number of predicted S-palmitoylated proteins with good experimental support (high FC, Supplementary Figure S1), it can be assumed that the false-positive rate is very high, unless experimental methods fail to identify a very large fraction of S-palmitoylated proteins.

### Palmitoylated proteins in S2R+ cells are enriched for subcellular membrane compartments

We performed a Gene Ontology (GO) term enrichment analysis to obtain a functional profile of the S2R+ cells palmitoylome (most significant terms in Figure 2, full list in Supplementary Table S4). In the cellular component category, “endomembrane system” and “intrinsic component of the membrane” were the most significantly overrepresented terms, followed by terms that refer to particular endomembrane system components such as the plasma membrane, Golgi apparatus, endoplasmic reticulum, cytoplasmic vesicles, or organelle membranes in general, but excluding the nuclear membrane and mitochondria. This is in agreement with the known common compartments of S-palmitoylated proteins (membrane compartments in general and the ER/Golgi to cytoplasmic membrane system in particular) [12] and the recently reported function of protein sorting for export from the Golgi apparatus [7].

**Figure 2:**
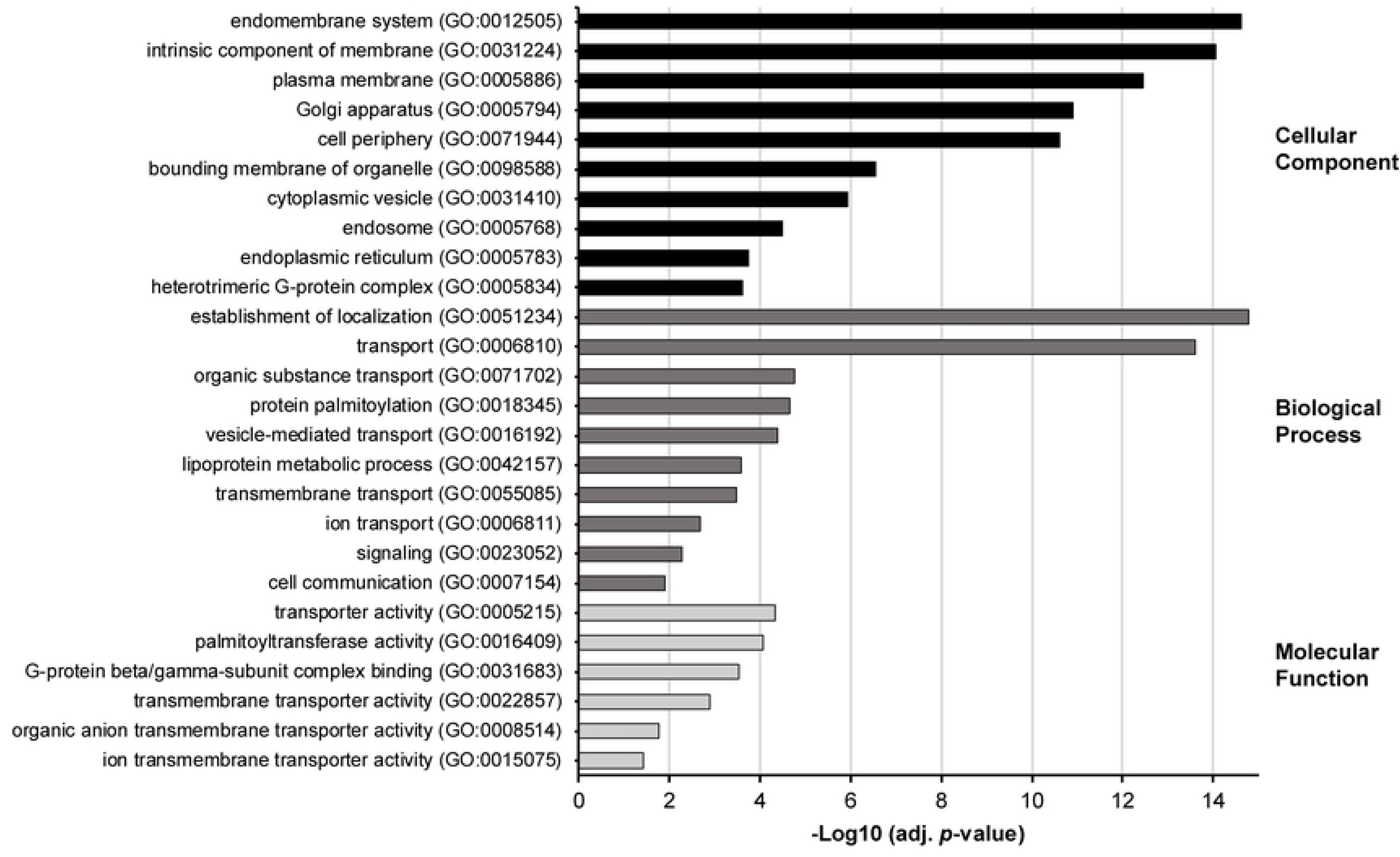
Most significantly associated GO terms with the Drosophila S2R+ cell palmitoylome proteins. Each bar represents the enrichment of a GO term in the dataset and they are separated in three main categories: cellular component (black), biological process (dark grey) and molecular function (light grey). These terms were manually selected to be the most representative and avoiding redundancy. Full list can be found in Supplementary Table S4.

Regarding function, the enrichment in the biological process and molecular function ontologies revealed that the palmitoylome of S2R+ cells mostly comprises proteins with localization and transporter activities, such as proteins involved in vesicle-mediated transport (SNARE proteins Bet1, Sec20 or Snap24), or transporter proteins that directly enable the movement of ions and molecules across membranes (Indy, JhI-21 or Nh3). Additional enriched functions include cell communication and signaling, particularly represented by the presence of the heterotrimetric G-proteins Galphai, Galphao, Galphas, Galphaq and cta (GO cellular component: heterotrimetic G-protein complex, and GO molecular functions: G-protein beta/gamma-subunit complex binding). Cycles of S-palmitoylation and de-acylation have previously been shown to regulate G protein-coupled receptor (GPCR) signaling activation and deactivation in a mammalian cell line [45] in a manner akin to the S-palmitoylation-dependent regulation of H- and N-Ras-subcellular localization and signaling [46]. The biological process “protein S-palmitoylation” and the molecular function “protein-cysteine S-palmitoylatransferase activity” were also enriched and of the 22 Drosophila DHHC-PATs, we found seven in the acyl-RAC dataset (Dnz1, GABPI, CG1407, CG5196, CG8314, CG34449/dZDHHC8 and dHip14) (Supplementary Data File SDF1 and Supplementary Figure S7).

Our data yielded the expected GO-term enrichments of S-palmitoylated proteins: membranes, endosomal/Golgi localization and transport and S-palmitoylation. The dynamic S-palmitoylation of GPCR signaling proteins described in mammals might also be important in insects. This is interesting, as the GPCR repertoire of invertebrates and vertebrates varies considerably, suggesting an origin of G-protein S-palmitoylation prior to the formation of the major GPCR superfamilies found in mammals [47].

### Optimization of experimental conditions for the use of BioID in Drosophila S2R+ cells

With the palmitoylome of S2R+ cells at hand, our next aim was to establish a method to identify interaction partners of DHHC-PATs. We used our S2R+ cell line as a starting point and explored the use of DHHC-PAT-BioID in combination with UAS/GAL4-dependent cell-specific expression in the fruit fly [48] as a means to acquire cell-specific DHHC-PAT interactomes.

To date, BioID has not been successfully used in Drosophila. Originally, BioID was developed for organisms adapted to 37 °C. Drosophila does not tolerate temperatures above 32 °C [49] and Drosophila S2R+ cells also died quickly after a shift in temperature from 25 °C to 37 °C (Supplementary Figure S2). Hence, neither flies nor Drosophila cells can be cultured at the temperature that BioID or BioID2 are usually used at (37 °C). Therefore, we carried out pilot experiments to determine the optimal conditions for the use of BioID and BioID2 in S2R+ cells. myc-BioID was successfully expressed using the pMt-Gal4/UASt system and resulted in low protein biotinylation activity in medium that was not supplemented with biotin (Figure 3A). BioID activity was markedly increased in cells grown in medium supplemented with 50 μM biotin, while 300 μM biotin did not further increase biotinylated protein abundance (Figure 3A). Cells grown at 30 °C showed slightly increased myc-BioID expression and 2-fold higher biotinylation activity compared to cells grown at 25 °C (Figure 3A and Supplementary Figure S3). These conditions were then used to compare the activities of BioID and BioID2. Cells harvested 24, 48 and 72 h after induction of expression showed comparable levels of expression of the biotin ligases and protein biotinylation increased with longer incubation time (Figure 3B). Relative quantification of biotinylation signals showed that in the presence of 50 μM biotin, BioID had higher activity compared to BioID2 (Figure 3C). This is in contrast to the superior performance of BioID2 in mammalian systems [50]. Therefore, we proceeded to use BioID under the optimized conditions determined above (i.e. 30 °C, 50 μM biotin and incubation times of 48 h) in all subsequent experiments with the S2R+ cell line. Moreover, BioID is also feasible in larvae and adult flies (Supplementary Figure S4 and S5). When using strong drivers or ubiquitous expression, myc-BioID is efficiently expressed, allowing the use of whole-animal lysates. However, in low-abundant cell types BioID did not yield enough biotinylated proteins to be efficiently detected in whole-animal homogenates. In that case, enrichment of body parts abundant in the target cell type (e.g. adult heads for neurons Supplementary Figure S4D) or use of TurboID [51], a recently bioengineered high-efficiency variant of BioID, may be recommended.

**Figure 3:**
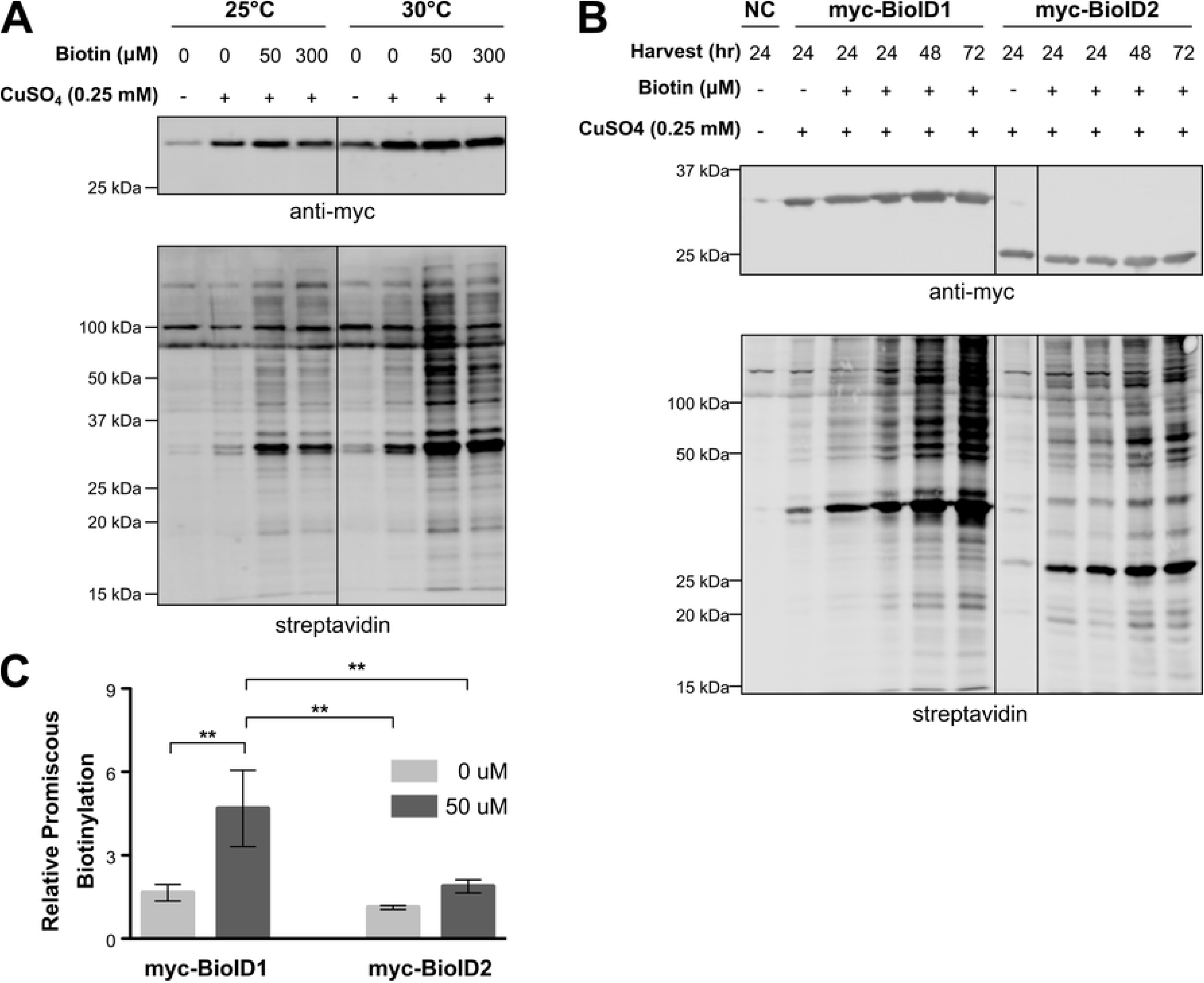
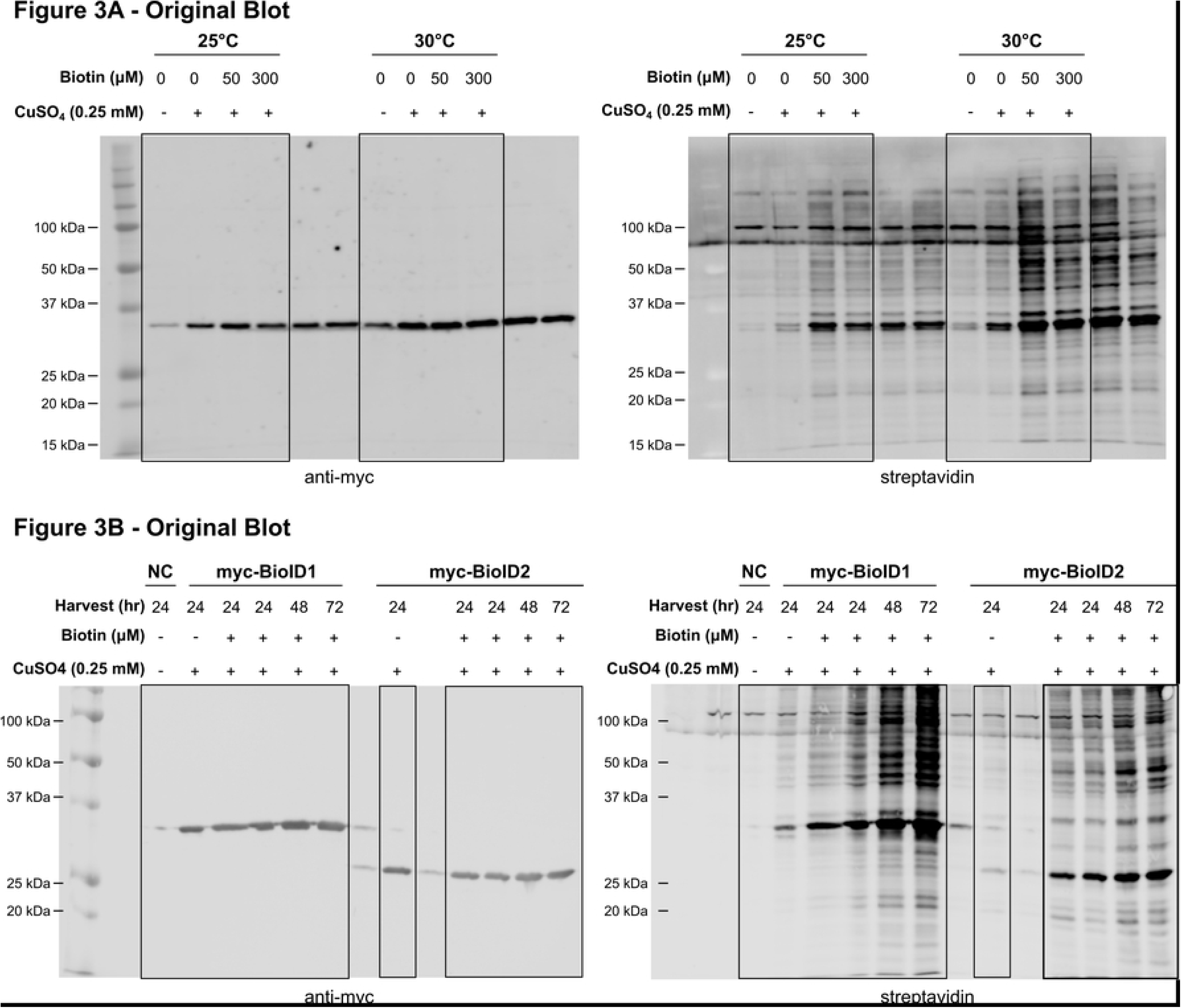
Establishment of BioID in *Drosophila* S2R+ cells. (A) Cells expressing soluble myc-BioID were incubated either at 25 or 30°C with medium containing indicated biotin concentration (0-300 uM) and harvested 24 hours post induction. (B) Overexpression of myc-BioID and myc-BioID2 in S2R+ cells incubated at 30°C at different time points (24, 48 and 72 hours) post-induction. (A, B) Western blots representative of cells expressing myc-BioID or myc-BioID2 using the Gal4/UASt system and induced with CuSO_4_ 0.25 mM. myc-BioID was detected using an anti-myc primary antibody (upper panels) and biotin was detected using a streptavidin probe (lower panels) on the same blot membrane using two different fluorophores on a Li-Cor Odyssey. (C) Quantification of promiscuous biotinylation relative to BioID or BioID2 overexpression in S2R+ cells 48 hours after induction and grown in medium in absence (0 uM) or supplemented with 50 uM biotin. Biotinylation signals are normalized to myc-BioID or myc-BioID2 expression levels respectively. Results from three independent experiments are shown as means +/- standard deviations. Asterisks indicate statistically significant differences in relative biotinylation (*, P<0.05; **P<0.01) according to one-way ANOVA with Bonferroni’s multiple comparison test.

### DHHC-BioID is specific enough to identify DHHC-PAT substrates

Having established BioID in Drosophila cells and flies, we proceeded to test whether DHHC-PAT-BioID constituted a feasible method to identify client proteins of DHHC-PATs. To this end we chose the well-characterized Drosophila DHHC-PAT Huntingtin-interacting protein 14 (dHip14). dHip14 is the ortholog of the mammalian HIP14 protein with which it shares the client proteins cysteine string protein (CSP) and Snap25 [21, 22]. In addition, HIP14 is the only known DHHC-PAT for which a conserved recognition motif in its client proteins has been identified (-[VIAP]-[VIT]-X-X-Q-P-) [52]. Importantly, the mutation of a proline to an alanine residue prevents the interaction of client proteins with HIP14 and thereby their S-palmitoylation [52].

To study the specificity of DHHC-BioID, we co-overexpressed dHip14-BioID or two unrelated DHHC-PATs, CG5620-BioID and CG6618-BioID, with N-terminally FLAG-tag-fused wt Snap25 (FLAG-Snap25) or the proline mutant (FLAG-Snap25*). Using Western blot and streptavidin probes to detect protein biotinylation we found a considerable increase in biotinylation of a protein with the relative molecular mass of dSnap25 in cells expressing dHip14-BioID and FLAG-dSnap25, but not in cells expressing dHip14-BioID and FLAG-dSnap25* or in cells expressing any of the other two DHHC-PAT-BioID fusion proteins in combination with FLAG-dSnap25 (Figure 4A, 4B and 4C for quantified data). Increased biotinylation was taken as evidence of successful DHHC-PAT-BioID expression and activity, while anti-FLAG antibodies were used to identify FLAG-tagged proteins and to confirm comparable expression levels of FLAG-Snap25 and FLAG-Snap25*.

**Figure 4:**
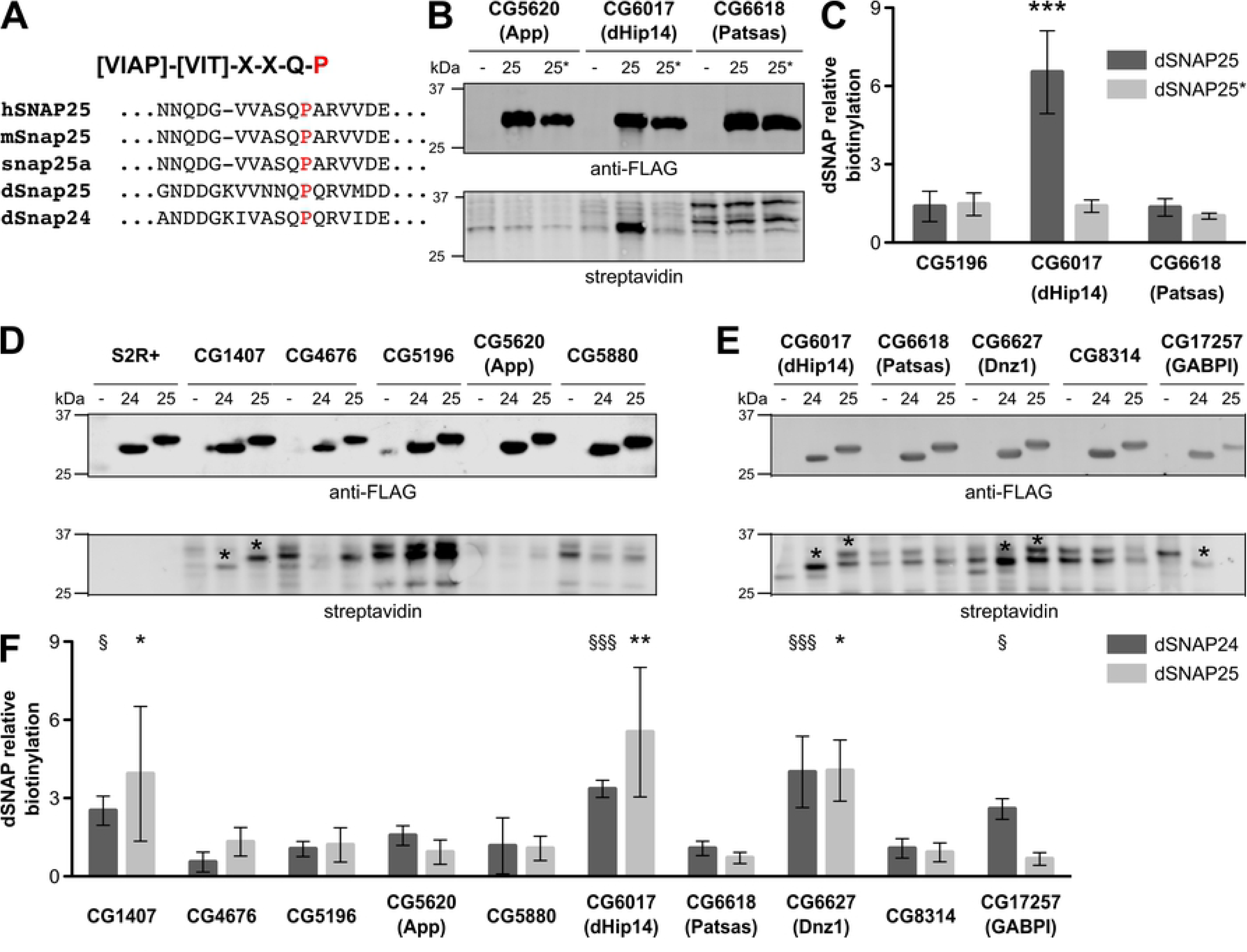
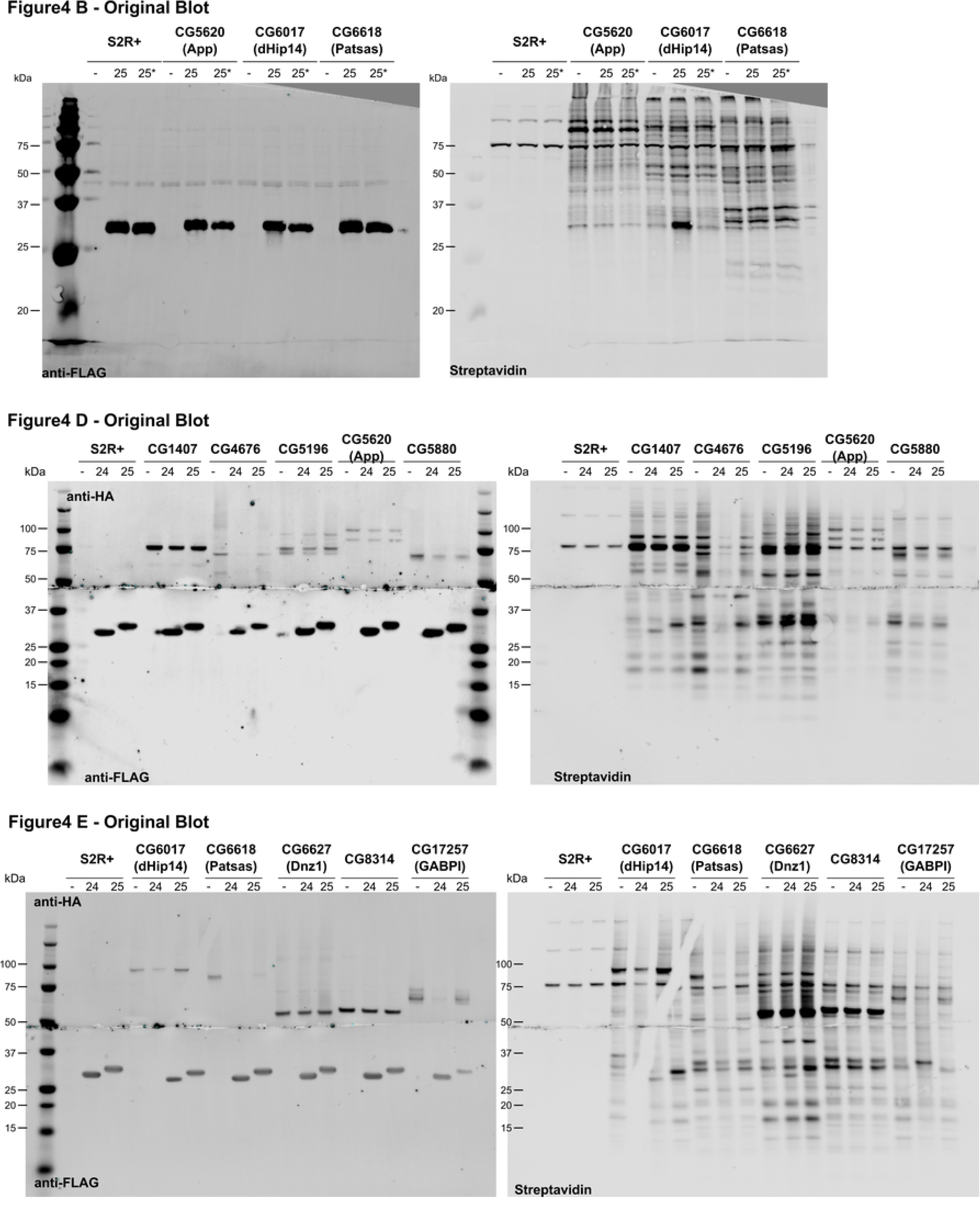
Validation of DHHC-BioID system using a known interaction. (A) Alignment of several orthologues and paralogues of SNAP25 in the region flanking dHip recognition motif ([52], grey box). Red: conserved Proline essential for the interaction with Hip14 and mutated to Alanine in dSNAP25* (P125A, 25*). (B) Western Blot representative of co-overexpression of CG6017 (dHip14, used as positive control) or CG5620 (App) or CG6618 (Patsas) as BioID fusion proteins with either FLAG-dSNAp25 wt (25) or FLAG-dSNAP25 P125A (25*). Note that biotinylation signal of the band relative to FLAG-SNAP25 is not present in the mutant (25*), indicative of specific interaction with the enzyme fused with BioID, as quantified in (C). (D, E) Western Blots representative of co-overexpression of FLAG-dSNAP24 (24) or dSNAP25 (25) with a subset of 10 different DHHC-BioID. Relative biotinylation of the substrates was quantified in (F). (B, D, E) Representative blots are shown. FLAG-dSNAP24/25/25* were detected using an anti-FLAG primary antibody (upper panel) and biotin was detected using a Streptavidin probe (lower panel). Biotinylation and FLAG-tagged proteins were detected on the same blot membrane using two different fluorophores on a Li-Cor Odyssey. (C, F) Quantification of dSNAP relative biotinylation signal, normalized by the signal of the respective dSNAP in wt S2R+ cells, was estimated according to material and methods. Results from three independent experiments are shown as means +/- standard deviations. Asterisks (*) or section signs (§) indicate statistically significant differences in relative biotinylation of dSNAP25 or dSNAP24 respectively in the negative control (S2R+) using a one-way ANOVA with Dunnett’s multiple comparison test (* and § P<0.05; ** and §§ P<0.01).

To further explore the specificity of the system, we extended the analysis to 10 selected DHHC-PATs are localized in various organelles. We excluded 12 DHHC-PATs with highly testis- or ovary-specific expression patterns [14, 53]. Thus, DHHC-BioID constructs were generated for, dHip14, CG8314, CG5196, CG5880, CG6618 (Patsas), CG5620 (app), CG17257 (GABPI), CG1407, CG4676 and CG6627 (Dnz1) (Supplementary Figure S6) and co-overexpressed either with FLAG-dSnap24 or FLAG-dSnap25. Expression of the DHHC-PAT-BioID fusion proteins alone served as the respective negative control to judge the pattern of protein biotinylation of each DHHC-PAT-BioID construct when the FLAG-tagged client proteins were not co-overexpressed (Figure 4D, 4E). dSnap24 has not been described as a dHip14-BioID substrate, but it shares a high degree of sequence similarity with dSnap25, including the Hip14-interaction motif (Figure 4A). Quantification of signals at the respective molecular masses of dSnap24 and dSnap25 revealed that three DHHC-PATs resulted in consistently increased biotinylation (CG1407, Hip14 and Dnz1) (Figure 4D - F). We also observed a reproducible interaction of the non-enzymatic DHHC-PAT GabPI with dSnap24.

From these results we conclude that DHHC-PAT-BioID is specific enough to recognize the interaction of dHip14 and dSnap25 even in overexpression conditions, which usually have the tendency of enhancing non-specific interactions.

### dHip14-BioID is sensitive enough to detect endogenous dSnap24 and dCSP

Finally, we wanted to know whether DHHC-PAT-BioID was sensitive enough to detect endogenous levels of DHHC-PAT interaction partners. To do this, we expressed dHip14-BioID in S2R+ cells, then affinity purified the biotinylated proteins with streptavidin beads and identified these proteins by mass spectrometry. S2R+ cells exposed to the same treatment but not expressing DHHC-BioID were used as a negative control. We found dSnap24 and dCSP among the proteins recovered from dHip14-BioID expressing S2R+ cells (Supplementary Data File SDF2). dSnap24 was robustly identified in all three dHip14-BioID samples and absent in two out of three control samples, while dCSP was only detected in one of three dHip14-BioID samples. The lower recovery of dCSP was in agreement with our co-overexpression experiments, in which biotinylation of dCSP co-overexpressed with dHip14-BioID was also markedly weaker than biotinylation of dSnap24 or dSnap25 (Supplementary Figure S6), even if expressed at similar levels). Since dCsp and dSnap24 have 15 and 13 lysine residues each, the lower degree of biotinylation of dCsp is not due to a lack of lysine residues. In addition, lysine residues are hydrophilic and are usually exposed on a proteins surface and they should therefore be accessible biotinylation. A reason why biotinylation of dCsp may be less efficient could be phosphorylation. In mammals, CSP-alpha can be phosphorylated in a lysine-rich region (Patel et al., 2016), and it is likely that dCsp can also be phosphorylated. Hence, phosphorylation of dCsp could reduce its propensity to be biotinylated. Other factors that may change level of biotinylation may be the duration of an interaction between a client protein and a DHHC-PAT (the longer the interaction, the more likely a protein is biotinylated) and how many times the proteins interact (e.g. proteins that repeatedly undergo palmitoylation and de-palmitoylation have higher chances to get biotinylated than proteins that interact only once with a DHHC-PAT during their life-cycle).

Nevertheless, in this initial dHip14-BioID proteomic dataset, we also found several proteins (e.g. Golgin-245, Golgin-84, Sec24AB, Sec24CD and Ergic-53) (Supplementary Data File SDF2) associated with the Golgi apparatus location of dHip14 [14] and vesicle transport. Whether or not they are valid client proteins or interaction partners cannot be judged at this point, as other DHHC-BioID datasets would be required to estimate specificity.

Taken together, the recovery of the endogenous dHip14 client proteins, dSnap24 and dCSP shows that DHHC-PAT-BioID is sensitive enough to detect endogenous interaction partners of DHHC-PATs.

### Dnz1 does not rescue Hip14 mutant flies

As we retrieved Dnz1 as a novel interaction partner of dCSP, dSnap24 and dSnap25 (Figure 4 and Supplementary Figure S6), we asked whether this interaction was Hip14-independent. In dHip14 mutant flies, failure of dCSP-S-palmitoylation and hence membrane association has been identified as the cause of lethality, as expression of dCSP fused to the transmembrane domain of neuronal synaptobrevin rescues pharate adult lethality of dHip14-mutant flies [22]. We thus reasoned that if Dnz1 interacts with and palmitoylates dCSP in a dHip14-independent manner, overexpression of *Dnz1* in neurons should rescue pharate lethality of *dHip14* mutant flies.

To test this, we crossed mutant dHip14 flies carrying *UAS-Dnz1* or *UAS-dHip14-BioID* (*w^-^; If/CyO; UAS-Dnz1-3xHA, dHip14^ex12^/TM6B, Tb, Hu* or respectively *w^-^; UAS-dHip14-BioID/CyO; dHip14^2^/TM6B, Tb, Hu*) with mutant *dHip14* flies carrying the *elavGAL4* driver for neuronal expression (*w^-^; elavGAL4/CyO; Hip14^2^/TM6B, Tb, Hu*). Flies cultured at 25 °C were rescued by neuronal expression of dHip14-BioID but not by neuronal expression of *Dnz1* (Table 1). These results show that even though Dnz1 and dHip14 both interact with Snap24, dHip14 is required for S-palmitoylation of dCSP. As Dnz1 and dHip14 reside in different subcellular compartments (endoplasmic reticulum and Golgi apparatus, respectively [14]), it is likely that they don’t have complementary functions. However, the exact function of Dnz1, in the context of its interaction with dCSP, is not clear. At this stage, our results only allow us to draw the conclusion that Dnz1 on its own is not able to palmitoylate dCSP in fly neuronal tissue.

**Table 1:**
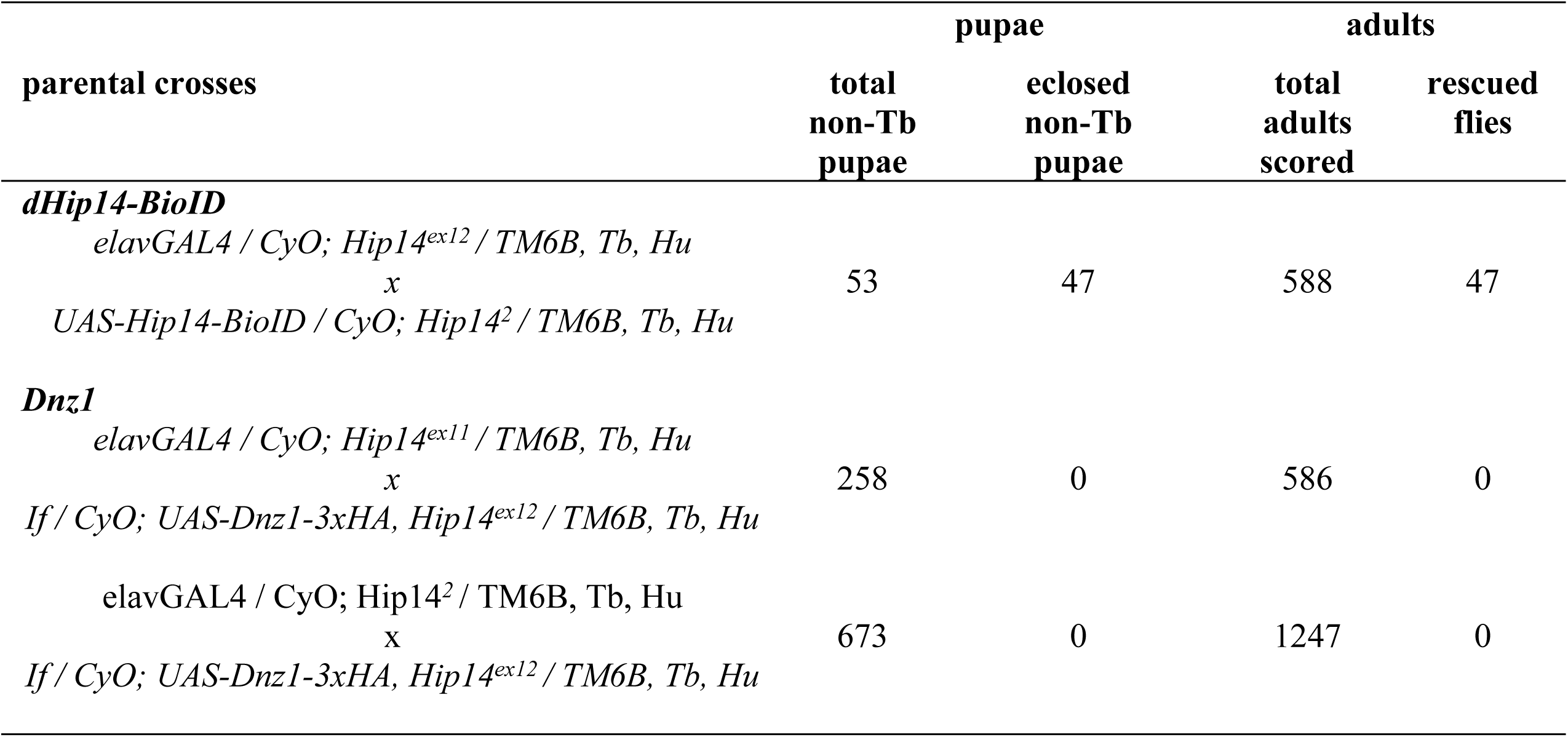
Rescue of *dHip14* mutant flies using neuronal expression of Dnz1 or Hip14-BioID

| parental crosses | total<br>non-Tb<br>pupae | eclosed<br>non-Tb<br>pupae | total<br>adults<br>scored | rescued<br>flies |
| --- | --- | --- | --- | --- |
| ***dHip14-BioID*** |  |  |  |  |
| *elavGAL4 / CyO; Hip14<sup>ex12</sup> / TM6B, Tb, Hu*<br>x<br>*UAS-Hip14-BioID / CyO; Hip14<sup>2</sup> / TM6B, Tb, Hu* | 53 | 47 | 588 | 47 |
| ***Dnz1*** |  |  |  |  |
| *elavGAL4 / CyO; Hip14<sup>ex11</sup> / TM6B, Tb, Hu*<br>x<br>*If / CyO; UAS-Dnz1-3xHA, Hip14<sup>ex12</sup> / TM6B, Tb, Hu* | 258 | 0 | 586 | 0 |
| *elavGAL4 / CyO; Hip14<sup>2</sup> / TM6B, Tb, Hu*<br>x<br>*If / CyO; UAS-Dnz1-3xHA, Hip14<sup>ex12</sup> / TM6B, Tb, Hu* | 673 | 0 | 1247 | 0 |

### DHHC-PAT-BioID identifies 180 putative DHHC-PAT interaction partners in S2R+ cells

With the proof of principle showing that DHHC-PAT-BioID is specific and sensitive enough to identify endogenous interaction partners of DHHC-PATs, we wanted to investigate the interactomes and putative client spectra of the aforementioned 10 DHHC-PATs in S2R+ cells. Thus, we carried out two independent experiments, each experiment consisting of three biological replicates per DHHC-PAT-BioID. In total, we identified 2162 proteins of which 487 proteins were enriched in at least one DHHC-PAT-BioID sample (FDR<0.1, FC>=2) compared with the negative control (S2R+ cells transfected with empty vector). These were subdivided in 193 proteins that reproducibly interacted with at least one common DHHC-PATs in both experiments (high-confidence DHHC-PAT interactors) and 294 non-reproducible putative interactors (low-confidence DHHC-PAT interactors) (Figure 5A). When compared to our previously established palmitoylome, we found that 25% of the palmitoylated proteins in S2R+ cells could be assigned to at least one DHHC-PAT; this suggests that DHHC-BioID is missing the interactions of three-quarters of palmitoylated proteins with their DHHC-PATs. Probable reasons could be: (1) only 10 out of the 13 non-testis-specific DHHC-PAT genes in Drosophila were investigated here; (2) of each of these, only one splice variant was used, whereas several genes are known to produce two or more differentially spliced products (Supplementary Figure S7); (3) DHHC-PAT interactions with their client proteins are transient and biotinylation efficiency of BioID used under the conditions described here may be limited to recovery of endogenous proteins that are abundant or undergo several cycles of S-palmitoylation/de-palmitoylation, increasing their chances to be labeled with a biotin.

**Figure 5:**
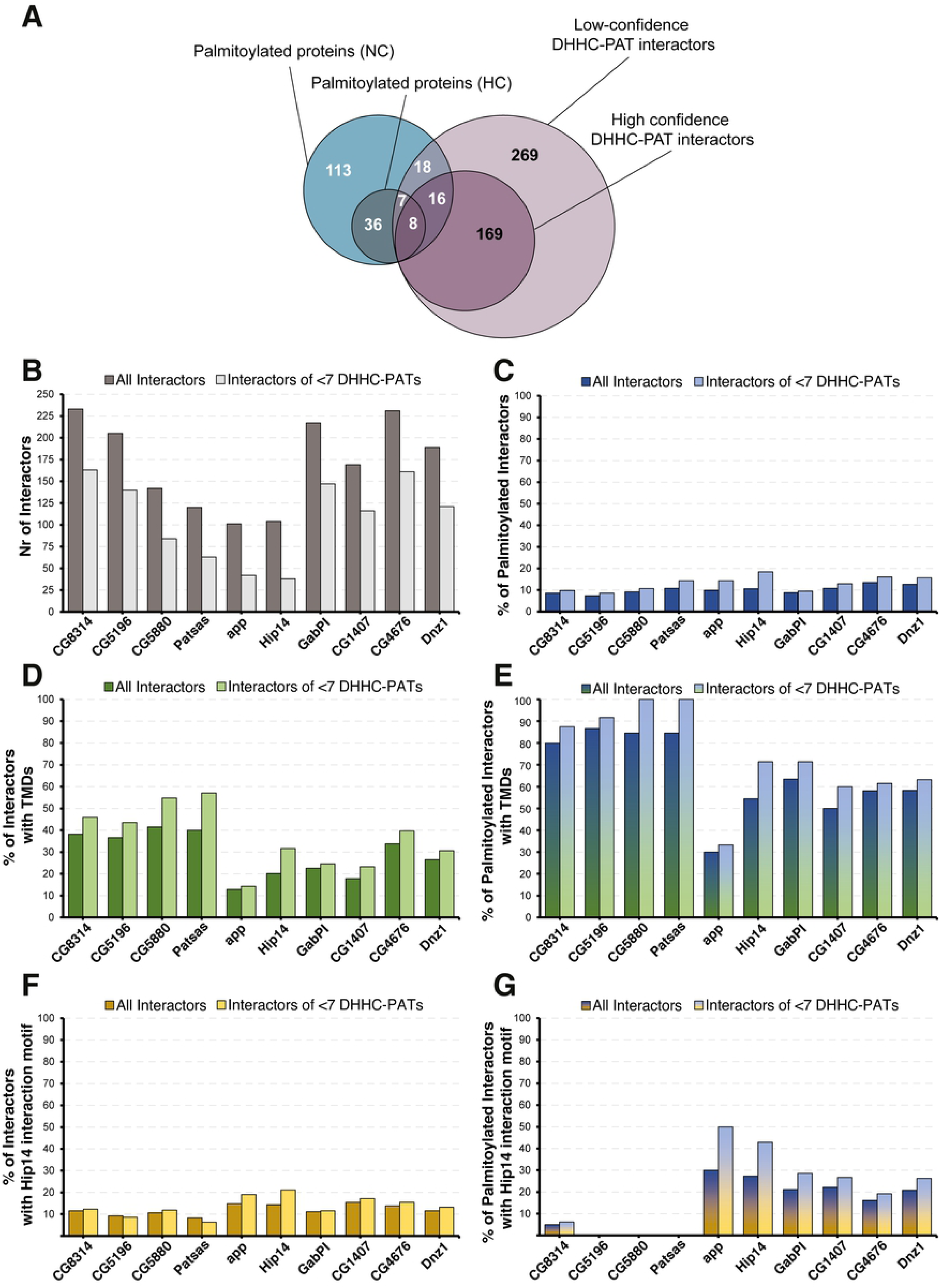
DHHC-PAT interactomes. (A) Venn diagram showing the overlap between the NC and HC fractions of the Drosophila palmitoylome of S2R+ cells and the DHHC-PAT-BioID interactome, which is also divided in low and high confidence interactors, depending on whether they were determined by one experiment alone or by both; (B) number of proteins (interactors) enriched by DHHC-PAT-BioID over the control. (C-G) percentage of DHHC-PAT-BioID interactors that (C) are palmitoylated, (D) are transmembrane proteins, (E) are palmitoylated transmembrane proteins, (F) contain the Hip14 interaction motif and (G) contain the Hip14 interaction motif and are palmitoylated. (B-G) ‘all interactors’ refers to proteins that are significantly enriched in the respective DHHC-PAT-BioID cell line over control cells; ‘<7 DHHC-PATs refers’ to interactors that are significantly enriched in the respective DHHC-PAT-BioID cell line over control cells and are not enriched in seven or more other DHHC-PAT-BioID cell lines.

Our main aim was to explore the potential of using proximity biotinylation for the identification of DHHC-PAT interactors. We find that this technique is suitable for the analysis of individual substrate candidates when ectopically expressing DHHC-BioID. This method may eventually permit the conclusive identification of specific DHHC-PAT interactions in Drosophila adult or larval tissues.

### DHHC-PAT-BioID interactors: S-palmitoylated substrates or neighboring proteins?

Since DHHC-PAT-BioID will not only recover client proteins that are S-palmitoylated by the DHHC-PAT under investigation, but also any other neighboring, accessory or co-localized protein within the 10 nm labeling distance of BioID [31], a significant challenge is to distinguish real client proteins of the DHHC-PAT from other interaction partners or neighbors that are not functionally relevant. Analyses of yeast and mammalian DHHC-PAT/client proteins have shown that most client proteins are unlikely to be S-palmitoylated by more than 4-6 different DHHC-PATs [54, 55]. In addition, in yeast, most of the seven DHHC-PATs can be knocked out without adverse effects, indicating that there is a large degree of redundancy between different DHHC-PATs [54]. To test whether the average number of DHHC-PATs that can palmitoylate the same protein could be derived from our dataset we checked whether the fraction of palmitoylated proteins that interact with a given number of DHHC-PATs peaked a specific number of DHHC-PATs. However, no such effect could be observed and no endogenous threshold could be derived from the dataset itself (Supplementary Figure S8). However, filtering for proteins that have a certain maximum number of DHHC-PAT- interactions will allow the identification of DHHC-PATs that actually have different substrate spectra. Of course, there might be protein/DHHC-PAT interactions that do occur with all DHHC-PATs, e.g. accessory proteins/enzymes, trafficking proteins, etc. So, to give a more complete picture, we present results from both approaches, i.e. a dataset to which no threshold for the number of DHHC-PAT interactions was applied, and a dataset in which a maximum of six DHHC-PAT-interactions (the maximum number of different DHHC-PATs known to palmitoylate the same protein in mammals [54, 55]) was allowed (Figure 5B-5G).

Little is currently known about different DHHC-PAT substrate specificities. According to two studies mammalian and yeast DHHC-PATs can be grouped according to their preference for soluble proteins and for integral membrane proteins [56, 57]. On the other hand, a high-throughput study in yeast reported that substrates of DHHC-PATs are possibly better grouped by the relative position of the cysteine residue that is S-palmitoylated (N-terminal, C-terminal or in respect to transmembrane domains) [54]. In our DHHC-PAT-interactome we found a certain bias of CG8314, CG5196, CG5880 and Patsas towards proteins with transmembrane domains (TMD); almost 40% of all of their interactors (both S-palmitoylated and non-S-palmitoylated proteins) had at least one TMD and around 80% of their S-palmitoylated interactors were TMD proteins (Figure 5D and 5E). For most other DHHC-PATs, the fraction of S-palmitoylated transmembrane substrates was around 50-60%, with the exception of CG6618 (approximated), which was markedly lower (30%).

To further investigate substrate specificities of DHHC-PATs, we checked for each DHHC-PAT in our study the fraction of proteins with the Hip14 interaction motif [52]. It ranged from 7% to 20% in different DHHC-PATs (Figure 5F). Furthermore, we observed an interesting correlation when considering S-palmitoylated substrates exclusively. Those DHHC-PATs that had the highest fraction of transmembrane proteins among their S-palmitoylated substrates also had the lowest fraction of S-palmitoylated substrates with the Hip14 interaction motif (0% in most cases) and vice versa, with the DHHC-PATs approximated and Hip14 showing the highest ratios (Figure 5E and 5G). These results and our co-expression experiments with Snap24, Snap25 and dCSP (Figure 4) show that even though Patsas and Hip14 both contain ankyrin repeats in their N-terminal cytoplasmic domains, only Hip14 interacts with its substrates via the Hip14 interaction motif. This is also in agreement with the observation that the two proteins are more distantly related than the two mammalian DHHC-PATs with ankyrin repeats (DHHC13/HIP14-like and ZDHHC17/HIP14) [14].

We also found that GABPI-PAT is among the DHHC-PATs with a broad range of interactors (Figure 5B), which may be indicating a role as a substrate-presenter rather than a palmitoyl-transferase [58].

## Conclusions

With our work we markedly increased the number of experimentally identified palmitoylated proteins in Drosophila and give a first detailed comparison between Drosophila and mammalian palmitoylomes and have highlighted certain functions, such as GPCR/G-protein signaling, where both groups are apparently similar in palmitoylation despite substantial differences elsewhere.

We also established DHHC-BioID as a novel method to identify proteins interacting with DHHC-PATs. This approach shows promise to also work in a tissue specific manner in Drosophila, and when combined with palmitoylome data increases the chance to identify client proteins for DHHC-PATs on the proteomic level.

## 4. Acknowledgments

We thank the Bloomington Stock Center, Maria Lind Karlberg (Uppsala University, Sweden), Norman Zielke (ZMBH, Heidelberg University, Germany), FlyORF (Zurick, Switzerland) and Steven Stowers (Montana State University, USA) for Drosophila stocks; Christopher Korey (College of Charleston, USA) for the Drosophila DHHC Gateway pENTR plasmid collection; the Sinning Group (BZH, Heidelberg, Germany) for plasmid pMT-GAL4 and the ZMBH proteomics facility (Heidelberg University, Germany) for mass-spectrometric analysis. This work was supported by DFG (Wi654/11-1). JC was supported by the German network for Bioinformatics Infrastructure (de.NBI) funded by the BMBF.

## Notes

### Competing Interest Statement

The authors have declared no competing interest.

